# A single locus carrying modified oogenesis genes underlies the switch to asexuality in Artemia brine shrimp

**DOI:** 10.64898/2026.04.06.716654

**Authors:** Marwan Elkrewi, Dominik Kopčak, Ariana Macon, Beatriz Vicoso

**Affiliations:** Institute of Science and Technology Austria (ISTA), Klosterneuburg 3400, Austria; Okinawa Institute of Science and Technology (OIST), 1919-1 Tancha, Onna-son, Kunigami-gun, Okinawa, Japan

**Keywords:** Meiosis, Asexual reproduction, parthenogenesis, single-nucleus RNAseq

## Abstract

Transitions from sexual to asexual reproduction are well-documented across different taxa. However, despite extensive efforts, the regulatory changes underlying the emergence of asexuality remain largely undiscovered in the majority of species studied. *Artemia* brine shrimp have multiple closely related sexual and obligate parthenogenetic lineages, making them a promising model for addressing this question. While earlier work suggested that asexuals use a modified meiosis, and inferred a likely role for the Z-chromosome in its transmission, no master regulator or genetic changes have been put forward as the root causes for the shift. Here, we generate single-nucleus RNAseq data of the female reproductive system of individuals from the Aibi lake population of *Artemia parthenogenetica* and its closely related obligate sexual species *Artemia sp. Kazakhstan*. We identify the germline cell clusters in the female reproductive system and perform differential expression analysis to infer substantial transcriptional differences at genes putatively involved in cell cycle and oocyte development between the meiotic cells of the two species. Additionally, we use whole-genome sequencing of 32 individuals from two backcrossing experiments to narrow down the genomic regions associated with the transmission of asexuality to an 8 megabase region of the Z chromosome. Within the identified regions, two adjacent genes with known functions in oogenesis, ITPR and USP8, show differential expression and genetic differentiation between sexuals and asexuals, making them promising candidate drivers of asexuality in this species.

**Significance statement:** While most animals reproduce sexually, many do not, and why and how these shifts occur remains an open question. This paper presents a systematic investigation of the molecular changes that underlie the transition from sexual to asexual reproduction in brine shrimp. We combine multiple computational and experimental approaches to look for differences between close sexual and asexual lineages. We find that a subset of meiotic germ cells is regulated differently in the two, and that two important oogenesis genes are the likely drivers of asexuality. This work is unique in providing an in-depth characterization of the combined genetic and regulatory changes underlying this key transition in reproductive modes.

## Introduction

The prevalence of sexual reproduction in eukaryotes suggests that it has advantages over asexual reproduction. Those advantages are thought to result from sexual populations being able to generate and maintain genetic diversity through meiosis and the exchange of genetic material between individuals, which in turn gives them a long term selective advantage (1). Asexuality, on the other hand, has often been described as a dead-end(2), and asexual lineages are typically short lived. However, many transitions from sexual to asexual reproduction (parthenogenesis) are observed in nature(3, 4). Asexual reproduction can be classified into apomixis (ameiotic reproduction) and automixis (modified meiosis). In apomixis, the individuals produced are genetic clones of the parent; automixis, on the other hand, can produce individuals identical or distinct from the parent, depending on the specific modification to the meiotic pathway and the presence or absence of recombination(5). While some of the initial triggers of asexuality are well understood (for instance hybridization or infection by a parasite/symbiont able to manipulate reproduction), what genetic and molecular changes encode the switch to asexual meiosis, and whether they involve simple steps or instead a complex remodelling of the meiotic program, are currently major gaps in our understanding of the dynamic evolution of reproductive systems. In particular, while a few loci and candidate genes have been identified in transitions from facultative/cyclical to obligatory parthenogenesis (6–8), how newly arisen parthenodes acquire the ability to both bypass the need for fertilization and achieve ploidy maintenance remains an open question (9).

Both apomixis and automixis are prevalent in arthropods, a large group where many transitions to asexuality have been reported(10). Brine shrimp (genus *Artemia*) are small crustaceans that live in hypersaline bodies of water throughout the world. While all new world species are sexual, many asexual lineages, both diploid and polyploid, have arisen recently in the Eurasian clade, and are known collectively as *Artemia parthenogenetica*. Classical cytogenetics studies were inconsistent on the mode of reproduction of the diploid parthenogenetic lineages of *Artemia*, as both gamete fusion and skipping of both meiotic divisions were suggested (11). However, recent evidence from both population genetics and cytology suggests that diploid parthenodes perform automixis by skipping the first meiotic division (11). Specifically, parthenogenetic oocytes do not arrest at metaphase I and do not divide after meiosis I (12). *Dmc1* and *Rad51*, two genes known to be involved in homologous recombination, were found to be upregulated in the sexual species *Artemia franciscana* compared to *A. parthenogenetica*, and knocking down *Dmc1* in this sexual species resulted in issues during homologous pairing (12). However, these inferences were potentially confounded by the phylogenetic distance between *A. franciscana*, which belongs to the American clade, and *A. parthenogenetica*, since the two lineages are estimated to have split around 40 million years ago (13–15).

On the other hand, comparisons of gene expression between various closely related sexual and asexual lineages in the Eurasian clade of *Artemia* using bulk RNA sequencing of ovarian tissue (16) did not show many consistent differences between the sexual and asexual lineages. While this could reflect the fact that only minor changes are needed to modify meiosis for asexuality, averaging gene expression across the tissue may hide subtle changes that occur in meiotic cells, since these only represent a small minority of cells in ovarian tissue. To circumvent those challenges, we generated single-nucleus RNAseq data from the female reproductive system of closely related diploid asexual (*A. parthenogenetica*, from a stock originally sampled from Aibi Lake, China) and sexual (*Artemia sp. Kazakhstan*) females. The single-nucleus resolution allowed us to capture different time points in the meiotic progression and identify differences between the sexual and asexual meiotic transcriptional pathways. Additionally, parthenogenetic lineages of *Artemia* are able to produce functional males at very low frequencies (17), which can mate with closely related sexual females to produce new asexual lineages, a phenomenon referred to as contagious parthenogenesis (18). We utilized the occurrence of three rare males to generate backcrosses to a sexual species (*Artemia sp. Kazakhstan*) for genetic mapping of the locus underlying the switch to asexuality. Finally, we combined the two approaches, along with variant calling and effect predictions, to narrow down two genes likely to encode the evolution of asexuality.

## Results

### snRNAseq provides insights into the (mis)regulation of meiotic programs in asexual germline cells

We performed single-nucleus RNA sequencing (snRNAseq) on the female reproductive system of the sexual *A. sp. Kazakhstan* and of asexual *A. parthenogenetica* (Aibi lake). Two replicates (comprising 25 females each) were obtained for each lineage. *Artemia sp. Kazakhstan* and the asexual lineages show very little genetic differentiation (16), making them ideal for direct comparisons. Genetic differentiation is also minimal with their close outgroup *Artemia sinica*, for which a high quality genome is available. We therefore mapped the reads of the two species to the published genome of *A. sinica* (after polishing it, see methods), and used the same pipeline as in (19) for preprocessing the snRNAseq data and clustering the inferred nuclei into putative cell types. The resulting UMAP is shown in Figure 1A (separate UMAP plots for the two lineages are provided in Sup. Fig. 1). The annotation of the different cell types was inferred through similarity in expression patterns with annotated cell types in a published dataset in the American species *Artemia franciscana* (using SAMap (20); Sup. Fig. 2 and 3) (19). All the recovered cell types mapped to a single cell type in *A. franciscana*, except AS_1 and AS_11, which aligned to both the *A. franciscana* Tracheal cells and Follicle cells (Sup. Fig. 2), and were each assigned to the cell type to which they had the highest alignment score. Two types of germ cells were recovered (germ cells A and B, Figure 1 A). Germ cells A correspond to early germ cells, as illustrated by their expression of the conserved early germline marker Orb, while germ cells B express the late germline marker Vasa. The two types of germ cells also express different sets of homologs of *Drosophila* meiosis genes identified in (19), confirming that they are made up of different stages of meiotic cells (Sup. Fig. 4 and 5).

**Figure 1:**
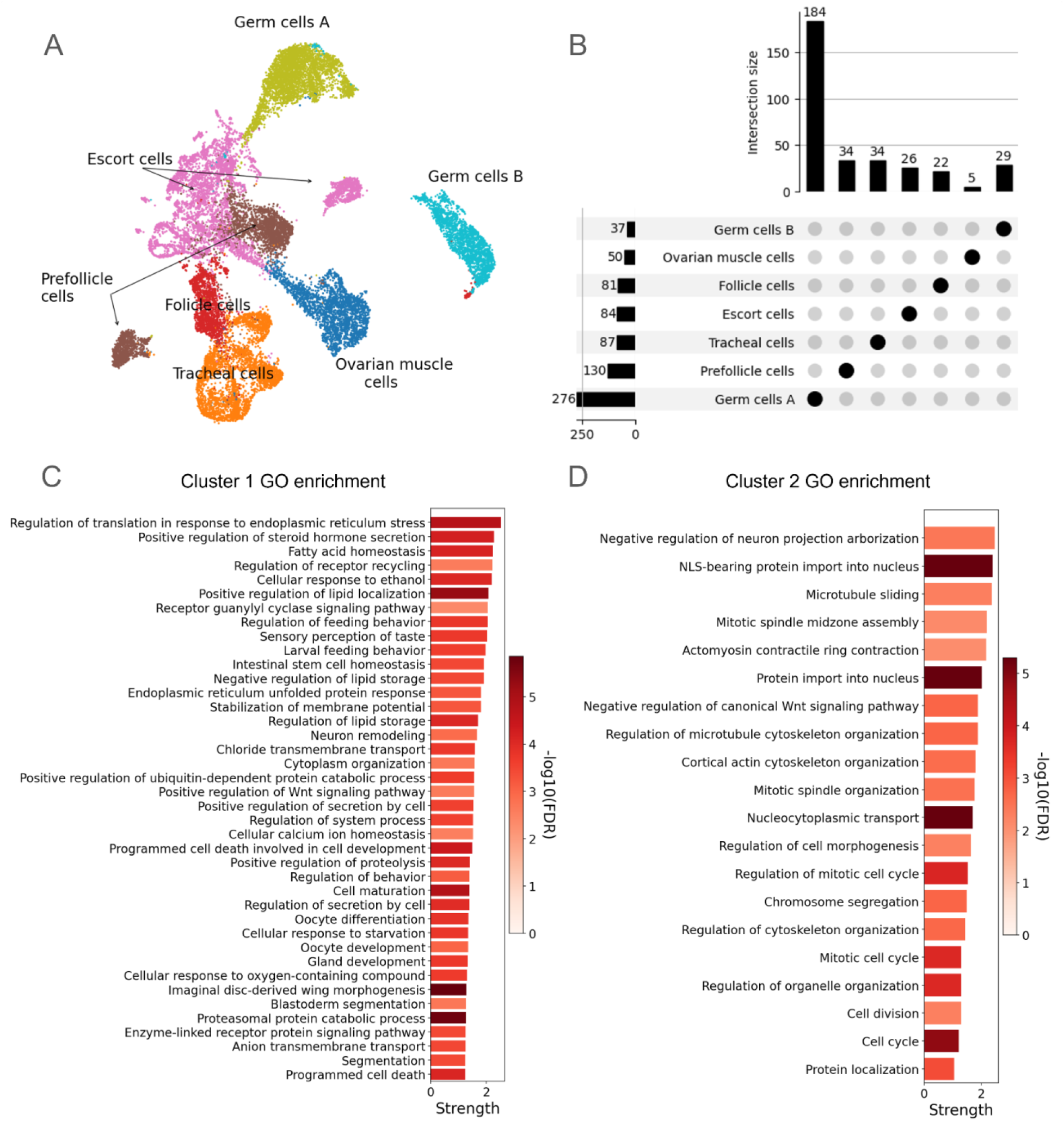
Early germ cells undergo most transcriptional changes in asexual *Artemia*. A) UMAP plot of the different cell types found in the reproductive system of both sexual and asexual lineages, annotated using the *A.franciscana* atlas. B) Total number of genes differentially expressed between sexual and asexual *Artemia* in both analyses for each cell type (horizontal bars), along with the number of differentially expressed genes that are uniquely found in each cell type (vertical bars). C) Gene ontology (GO) enrichment detected in the first cluster from the stringDB k-means clustering analysis. D) GO enrichment detected in the second cluster from the stringDB k-means clustering analysis.

**Figure 2:**
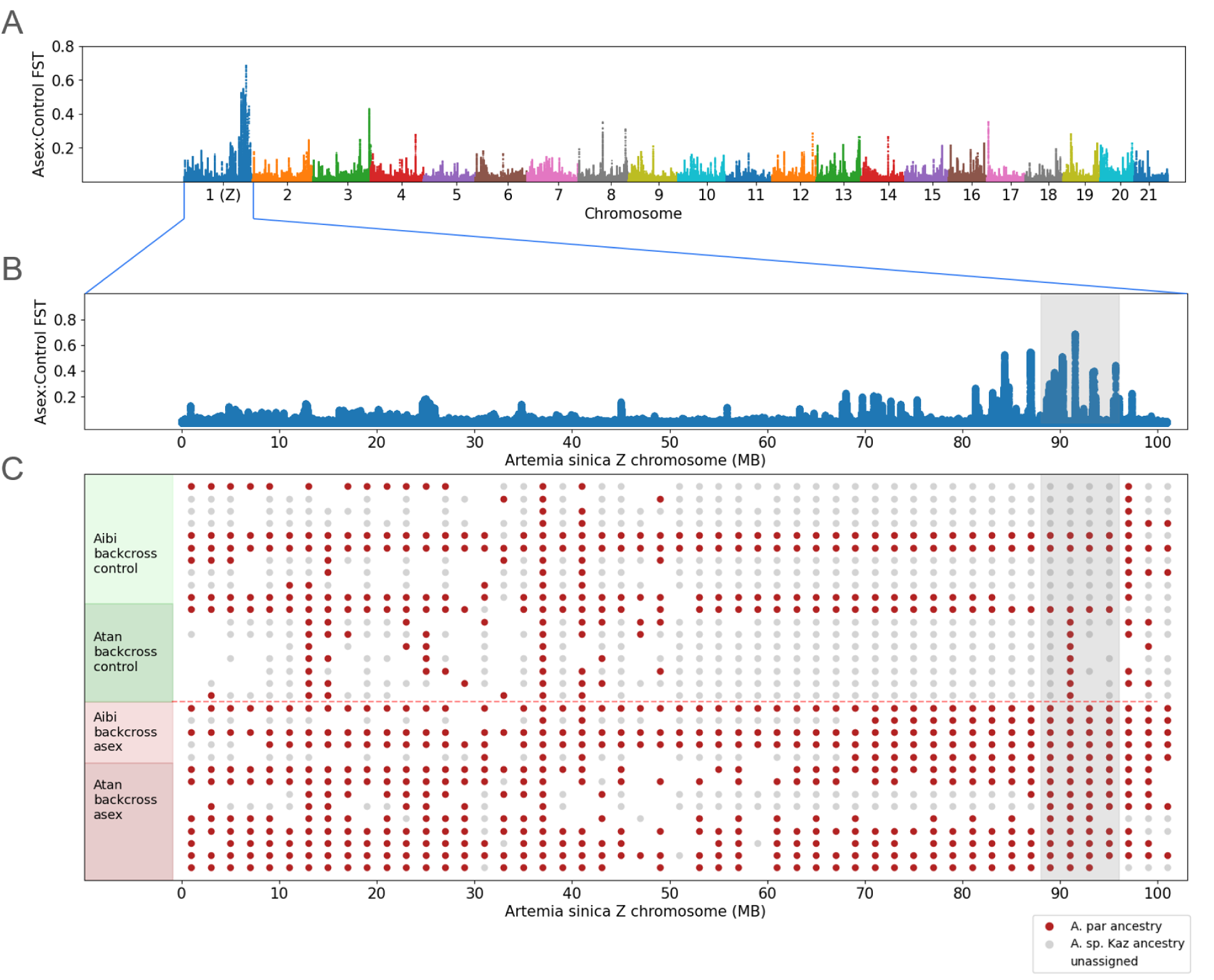
Genetic mapping yields a single locus on the Z chromosome underlying transmission of asexuality. A) FST between the asexual and control backcrossed F2 females along the genome (rolling median of FST estimates for 100 consecutive positions is shown) B) The same, but zooming in on the distal end of the Z chromosome. C) Inferred ancestry of 2MB windows along the Z chromosome (grey denotes *A. sp. Kazakhstan* ancestry only, red denotes *A. parthenogenetica* ancestry or both *A. sp. Kazakhstan* and *A. parthenogenetica* ancestry, white denotes unassigned). The region shaded in grey highlights windows with *A. parthenogenetica* ancestry in all the asexual individuals. Similar plots for other chromosomes are available in Sup. Fig. 6.

**Figure 3:**
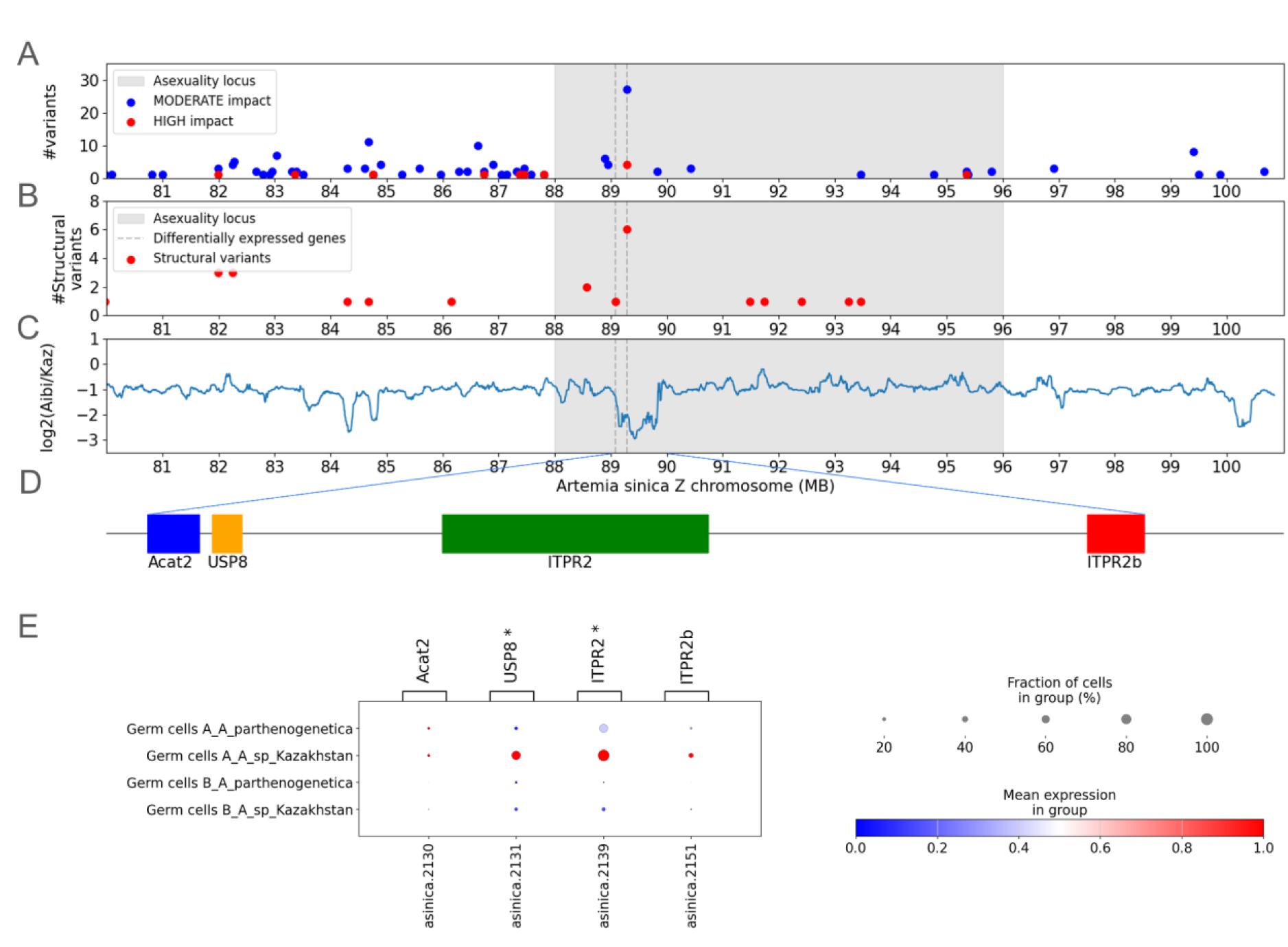
ITPR2 and USP8 show functional differentiation consistent with drivers of asexual reproduction. A) Number of short nucleotide variants per gene with high and medium predicted impact on protein function. B) Number of structural variants with high impact per gene detected from long DNA reads. C) Differences in genomic coverage between *A. sp Kazakhstan* and *A. parthenogenetica* (Aibi lake). The shaded area represents the inferred location of the asexuality locus, and vertical lines show the location of differentially expressed genes. D) A schematic of the four genes in the region between 89 and 90 MB. E) A dot plot depicting the expression of the four genes in the region between 89 and 90MB in the germ cells A and B.

**Figure 4:**
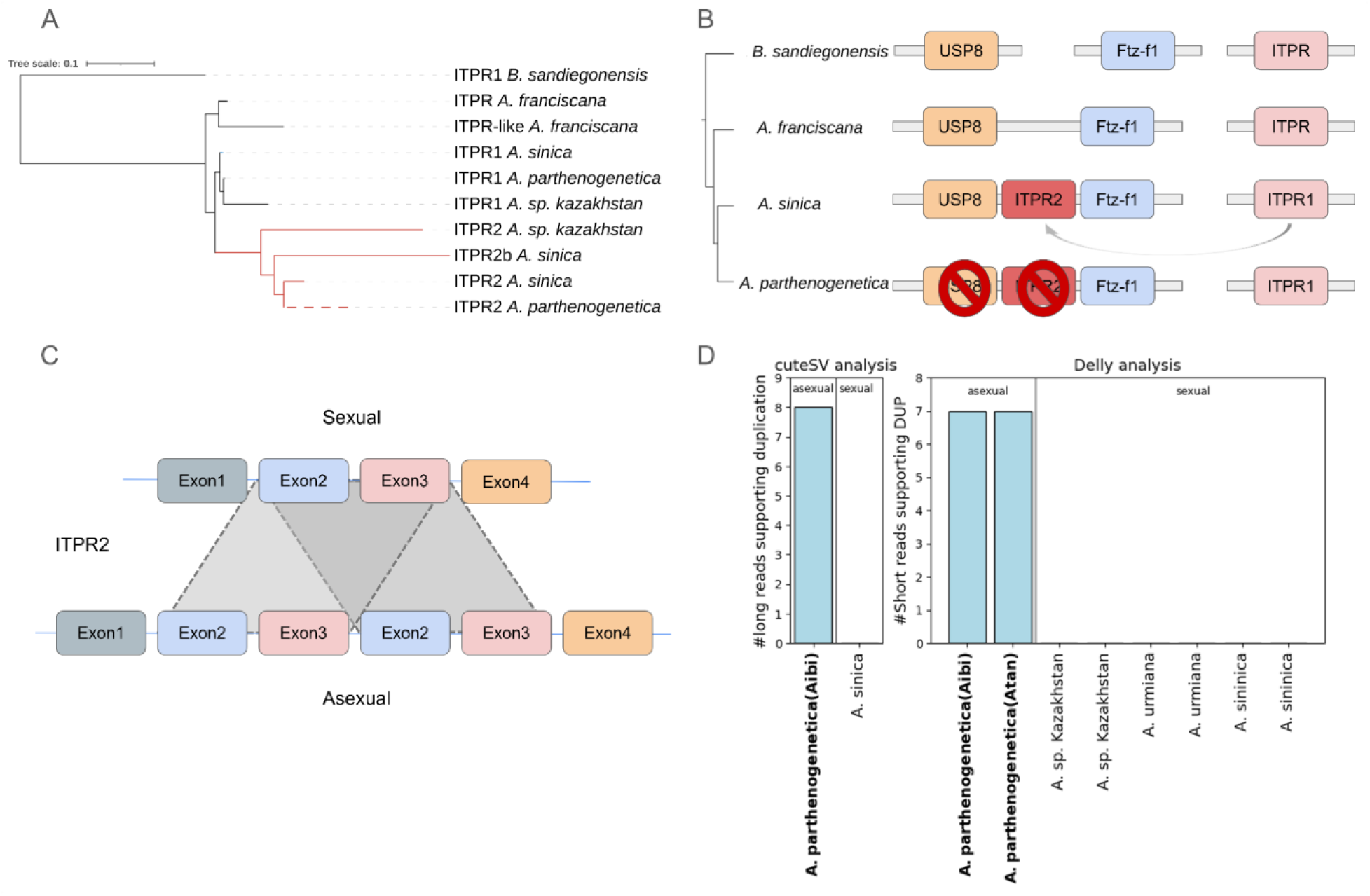
Evolution of the differentiated region of the asexuality locus. A) Phylogenetic tree of ITPR homologs across *Artemia* species shows that ITPR2 arose through a duplication in the Eurasian lineage. B) Schematic model for the evolutionary changes that lead to the transition to asexuality. C) Schematic of the duplication of the second and third exons detected in asexual *Artemia*. D) Number of long (CuteSV) and short (Delly analysis) reads supporting the duplication in the asexual and sexual lineages of *Artemia*.

**Figure 5:**
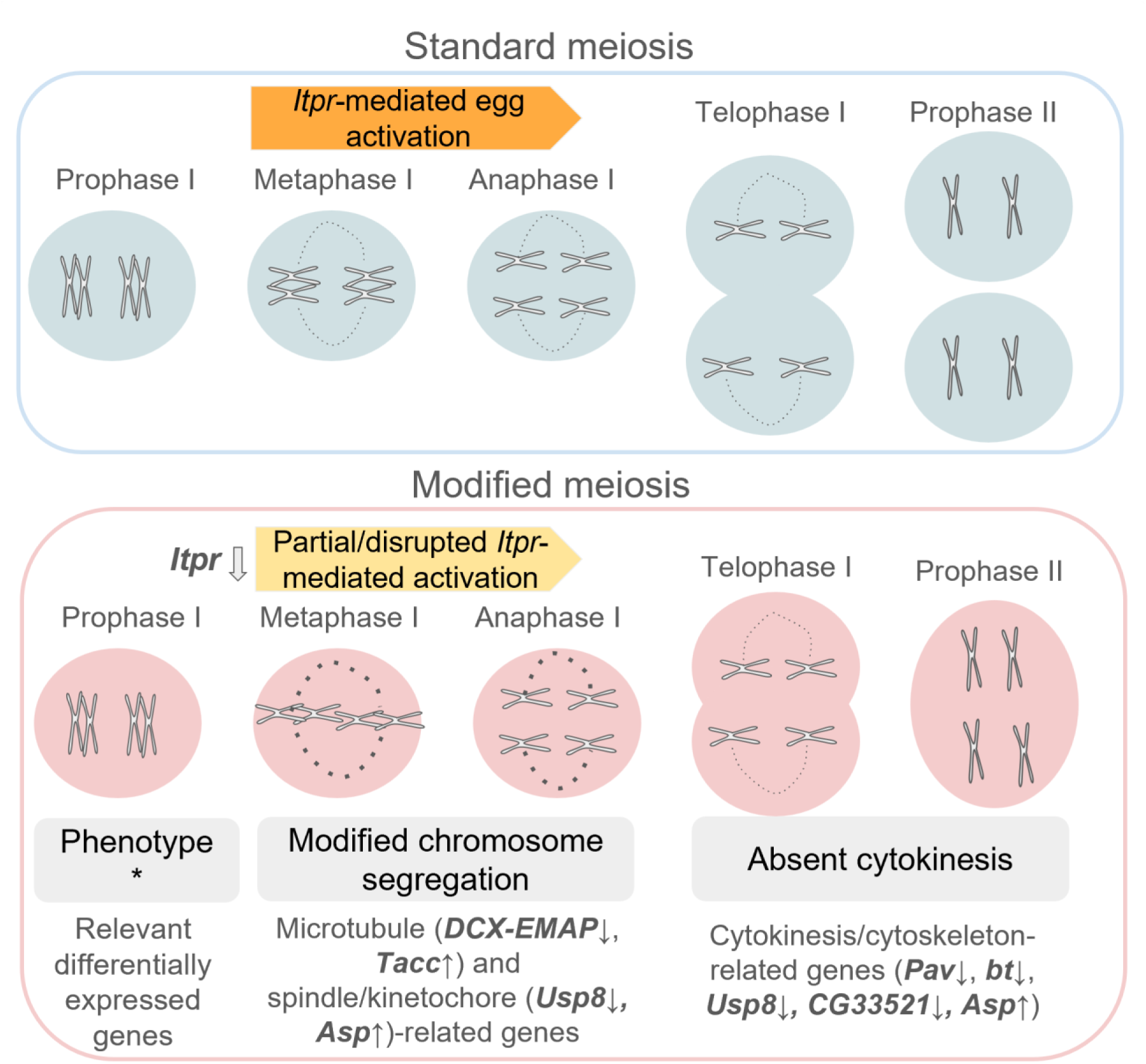
A model for the ITPR-based evolution of asexual meiosis. Gene names refer to Drosophila orthologues of top differentially expressed genes in germ cells A. Arrows next to the names show the direction of the change in expression in *A. parthenogenetica* relative to *A. sp. Kazahkstan*. * Cellular phenotypes from (12).

In order to identify genes that could be involved in the regulation of asexual meiosis, we performed differential expression analysis between the sexual and asexual lineages within each cell type using two complementary approaches: pseudo-bulking and DEseq2(21), and Seurat(22) Findmarkers and MAST(23) (see methods for details). We focused on genes that were significantly biased in the two analyses (p-adj <0.05 and fold change >2) (Figure 1 B). Germ cells A had the largest number of DE genes (276 genes, with 185 upregulated in the sexual species and 91 upregulated in the asexuals), suggesting that the transcriptional changes that lead to asexual meiosis occur at this early stage of germline development (Figure 1 B). Surprisingly, Germ cells B had only 37 DE genes, the lowest number of all cell types, suggesting a lack of transcriptional differences later in oogenesis. We performed a functional enrichment of genes differentially expressed in germ cells A with stringDB. (24). This analysis recovered no significant enrichment in protein-protein interactions; however, the network had a large number of significant gene ontology enrichments (Sup. Table 1). We used the k-means clustering algorithm on stringDB to check if the differentially expressed genes can be grouped into clusters that affect distinct pathways. With the default number of clusters (3), we recovered a cluster of 69 genes with GO terms related to oocyte development (Figure 1 C), a cluster of 33 genes enriched for cell cycle related terms (Figure 1 D), and a cluster of 2 genes with no GO enrichments. This is consistent with the observation that *A. parthenogenetica* has continuous oogenesis (12), which would require changes to allow meiosis to proceed without an arrest and oocytes to develop without fertilization.

### Genetic mapping identifies a single Z-linked asexuality locus

Previous genetic mapping using F2 females from a backcross of an *A. parthenogenetica* (Aibi lake stock) rare male and a sexual *A. sp. Kazakhstan* female (25) suggested a role for the Z chromosome in the transmission of asexuality. These hybrid backcrosses yield few individuals, of which only a small number of females reproduce asexually (5 in the previous study), strongly limiting their power. We therefore repeated the experiment using a cross between three rare males from *A. parthenogenetica* (Lake Atanasovsko stock) and *A. sp. Kazakhstan* females. Male offspring from the F1 were backcrossed to unmated (sexual) *A. sp. Kazakhstan* females. From the F2s, we obtained and sequenced 9 additional females which reproduced asexually, and 8 control females (which did not produce offspring after 3 months of isolation). We aligned the genomic sequencing data of the 32 individuals from the two experiments to the polished *A. sinica* genome assembly (25). We called SNPs and estimated the Asexual:Control F_ST_ for all biallelic SNPs to find genomic regions showing genetic differentiation between the two types of females, as expected of loci involved in asexuality. A smaller region (∼18 MB) showed elevated F_ST_ on the Z chromosome (Figure 2 A and B), compared to the previous analysis(25). We further characterized the patterns of inheritance of non-overlapping 2 megabase (MB) windows across the Z chromosome. In each F2 individual, windows were either assigned *A. parthenogenetica* ancestry (if 60% or more of their SNPs had the *A. parthenogenetica* allele), or *A. sp. Kazakhstan-only* ancestry (if 60% of their SNPs or more were homozygous for the *A. sp. Kazakhstan* allele). Since all F2 asexual females must necessarily have inherited their asexuality loci from their *A. parthenogenetica* grandfather, we searched for windows where 100% of the asexual F2 but fewer than 50% of the control females had *A. parthenogenetica* ancestry (the mendelian expectation if controls are not enriched for sexual females, a conservative approach; reducing this cut-off to 30% does not affect the main conclusions, see Sup. Fig. 6). This filtering recovered only 3 windows, all in the region between 88 and 96 MB on the Z chromosome (Figure 2 C). As no other genomic regions showed consistent patterns of segregation with asexuality (Sup. Fig. 7), our results suggest that there is a single asexuality locus within this ∼8 MB region of the Z chromosome. The region contains 50 protein coding genes (Sup. Table 2).

### A differentiated region of the asexuality locus contains two key oogenesis genes

Our ability to infer the boundaries of the asexuality locus (see previous section) is limited by the small number of individuals obtained in the crosses. However, any gene in this region that plays a causal role must also be functionally different between sexuals and asexuals, which should be reflected in differences in either their protein sequence or gene expression. Only 2 out of 50 protein coding genes in this region, located adjacently at around 89.0Mb, were among our differentially expressed genes in germ cells A (Figure 3 A): a homolog of *ITPR*, which encodes a calcium channel known to be involved in oocyte maturation in various species, and a homolog of *USP8* (a deubiquitinase known to be involved in endoreplication and germline cell division in *Drosophila*). Both were strongly underexpressed in asexual germ cells A compared to sexual germ cells (adjusted p-value<10^−12^). An additional partial copy of *ITPR* is found in the region, and also underexpressed in asexual germ cells A, but not significantly so. Both copies are also underexpressed in the ovaries of various asexual lineages compared to those of multiple sexual species (data from (16); Sup. Fig. 8). We name these copies *ITPR2* and *ITPR2b*, as an additional copy of this gene is found on chromosome 5 (*ITPR1*; its patterns of expression, which do not differ between sexuals and asexuals, are shown in Sup. Fig 8).

Two more lines of evidence support *ITPR2* as a prime candidate for the regulation of asexuality. We identified genetic variants that are likely to disrupt protein coding genes in the asexual lineages by mapping published genomic reads derived from *A. sp. Kazakhstan* and *A. parthenogenetica* females (one from Aibi lake and one from Lake Athanasovsko) to the *A. sinica* genome. We used SnpEff (26) to predict the effects of the variants found exclusively in *A. parthenogenetica*. Most of the SNP and structural variants with a predicted functional effect, including 4/5 variants with a predicted strong effect, were located on ITPR2. Only one additional gene located in the asexuality locus carried a predicted high impact variant: a homolog of the *Drosophila* circadian gene *Nocte* (annotated as *PRRC2A-like*), located at 95MB, which carried a frameshift mutation. Closer inspection of the alignments showed that this was a misaligned point mutation at a codon that is not conserved between *A. kazakhstan* and *A. sinica*, and which replaces a non-polar amino acid (proline) with another (leucine), arguing against a strong effect of the change. In addition to single-nucleotide polymorphisms, we detected large structural variants that overlapped genes in the asexual species from newly generated PromethION nanopore long DNA sequencing reads from an *A. parthenogenetica* female (Aibi lake) and from an *A. sinica* female. Structural variants specific to the asexual female were detected using cuteSV (27), and their impact predicted with VEP (28). Although this analysis is noisy due to the use of only two individuals, ITPR2 had the largest number of structural variants with predicted high impact (Figure 3 B, Sup. Table 3). Finally, a broader region from 89 to 90MB also showed differences in genomic coverage between A*. parthenogenetica* and *A. sp. Kazakhstan*, in agreement with it being a highly rearranged and/or diverged region between sexuals and asexuals. Even long reads failed to align across lineages in parts of *ITPR2*, making it difficult to infer the specific rearrangements at play, but supporting the presence of a highly diverged region (Sup. Fig. 9). A fourth gene, *Acat2*, was found in this region. However, it encodes an enzyme involved in cholesterol metabolism, and showed very low levels of expression in the germline, making it an unlikely candidate. Taken together, these results show that *USP8* and *ITPR2*, two adjacent genes on the asexuality locus whose homologs are known to play key roles in meiosis and oogenesis, show expression (and sequence in the case of *ITPR2*) patterns most consistent with a causal role in asexual meiosis.

### Acquisition and evolution of Itpr on the asexuality locus

To investigate the origin of the two copies of *ITPR* on the Z chromosome of *A. sinica*, we searched for homologs of Itpr in the genome of the American species *A. franciscana* (diverged from the Asian species ∼30 million years ago). We found two copies on chromosome 5 (annotated as *ITPR* and *ITPR-like*), and no copy on the Z chromosome, despite the presence of a syntenic region containing *USP8* and *Ftz-f1*, the two genes adjacent to *ITPR2* in Eurasian species. To explore the phylogenetic relationship of those two copies and those of other Artemia lineages, we produced a phylogenetic tree with *ITPR* protein sequences obtained from the genomes of all available *Artemia* and of their outgroup, the fairy shrimp *Branchinecta sandiegonensis* (Figure 4A). All Artemia species share the *ITPR* copy on chromosome 5, which does not show any differences in expression between sexual and asexual species (Sup. Figure 8). The tree further suggests that *ITPR2* resulted from a duplication of *ITPR* into the Z chromosome in the ancestor of the eurasian species. The two *A. franciscana* copies cluster together, suggesting the shorter *ITPR-like* is likely the result of an independent duplication event that occurred after the split from the eurasian species. The third copy in *A. sinica* (*ITPR2b*) clusters with the Z chromosome copy *ITPR2*, and it is not clear whether it represents an independent duplication event or an artifact of the assembly process.

Among the 6 structural variants (SV) that we detected in *ITPR2* using the long DNA reads (Sup. Table 3), we identified a ∼20,000 bp duplication that spans the second and third exons fully (Figure 4 A and B). This SV was independently detected exclusively in two asexual lineages using Delly (29) with genomic short reads (Figure 4 A). As duplications of exons are known to create frameshifts that result in premature stop codons and lower mRNA levels due to non-sense mediated decay, this is one possible mechanistic explanation for the observed difference in mRNA levels between the sexual and asexual species. Overall, our findings show that *ITPR* was duplicated in the ancestor of Eurasian *Artemia* into a locus already containing *USP8*. While the sexual species maintained a functional *ITPR2*, the asexual lineages acquired modifications that resulted in the downregulation of this duplicate and of its adjacent gene *USP8*, either concurrently or sequentially, to create the current asexuality locus (Figure 4 C).

## Discussion

### A simple genetic architecture of asexuality

The recent generation of extensive genomic and transcriptomic resources in clades containing sexual and asexual lineages has allowed the identification of a few loci and candidate genes for the shift (6–8), suggesting a relatively simple genetic architecture underlying this transition. In *Daphnia pulex*, for instance, a transposon insertion upstream of one of the three copies in the genome of the meiotic cohesin Rec8 and a frameshift mutation seem to be shared across the asexual lineages, making the locus a likely candidate for the transition to obligate asexuality (6). In the Cape Honeybee, a Hymenopteran-specific protein, with 3 non-synonymous variants and a putative role in chromosome segregation, co-segregates with asexuality and is downregulated in asexual bees (8). However, these studies were performed on species that had parthenogenesis as part of their life cycle (either as cyclical parthenogenesis or haplodiploidy, where males arise from unfertilized eggs), and how de novo shifts to asexuality are encoded is still unclear. One key step in this direction was the identification of differentially expressed genes in the facultative parthenode *Drosophila mercatorum* (the cell division kinase *Polo*, the desaturase *Desat2*, and the transcription factor *Myc*), which when mis-expressed in *D. melanogaster* led a subset of females to reproduce through parthenogenesis(30). However, modulation of (unrelated) control genes also yielded a small fraction of parthenodes, and some strains of *D. melanogaster* are known to produce some offspring asexually, suggesting that the setup for automixis is already present in this clade. Fewer changes may therefore be required than for de novo origination of parthenogenesis.

We previously detected an association between the distal end of the Z-chromosome and the ability to reproduce asexually in Artemia (25). However, only a small minority of backcrossed females produced offspring asexually in our crosses (5, versus 96 that did not). One possibility that could therefore not be excluded at the time was that multiple loci across the genome were required for asexual reproduction. Here, we were able to narrow down the required region to a single locus containing 50 genes. Of these, only a few showed discernible differences in sequence or expression between sexuals and asexuals, suggesting that even a novel transition to asexuality only strictly requires one or a few genes to be modified for parthenogenesis to be possible. It should be noted that our stringent differential expression filtering may miss subtle or inconsistent changes at other genes, but at least exclude a general regulatory remodeling of many of the 50 genes in the asexuality locus. Importantly, these hybrid parthenodes produce few offspring, and other genes elsewhere in the genome likely further contribute to fully functional parthenogenesis (but are not strictly necessary). The asexuality locus itself includes several homologs of genes that could potentially play an additional role in modified cell division, including ftz-f1 (known to be involved in mitotic cell cycle progression)(31), Centriolin (important for centrosome function and cell division)(32), and PP1 (involved in the regulation of mitotic chromosomal segregation as well as regulation of chromatin condensation during interphase)(33).

The presence of a single locus promoting asexuality may also help shed light on the complex relationship between sexuals and asexuals. Evidence from flow cytometry in addition to mitochondrial and nuclear DNA data suggested that all the *A. parthenogenetica* lineages arose within the last 80,000 years through recurrent hybridization of an ancestral asexual lineage (34). Our backcross experiment used *A. parthenogenetica* males from a different stock (originally collected from Lakes Atanasvosko in Bulgaria) than the one used in the published backcross experiment (originally from Aibi Lake in China). The fact that both yielded the same region on the Z-chromosome as the likely locus underlying asexuality (Figure 2) supports a common origin of all asexuals. The transmission of a single locus is also more likely than that of a polygenic trait, potentially contributing to the recurrent creation of new asexual lineages.

### Early germline cells show most differential expression, but no change in DMC1/RAD51

Cytological evidence suggested that the oocytes of *A. franciscana* and asexual *Artemia* show the most differences in chromosomal morphology during middle oogenesis, when the oocytes move to the oviducts (12). This is also the stage where differential expression of the meiotic recombination and double-stranded breaks genes, *DMC1* and *RAD51*, was observed between one asexual lineage and its distant sexual relative *A. franciscana* (12). Although neither of those genes show significant expression differences between the sexuals and asexuals in our analysis (Sup. Fig. 10), *DMC1* was strongly expressed in germ cells A, the earliest germ cells we detect and where we find the largest number of differentially expressed genes, but less expressed in later germ cells B (Figure 3 D). These patterns therefore suggest that while the two approaches agree on the developmental stage at which meiosis is modified, the differences in DMC1 and RAD51 that were previously detected may reflect general differences between the American and Eurasian lines that were compared. This is in line with the fact that the Eurasian branch that gave rise to asexuals lineages has much lower recombination rates than the American *A. franciscana*(35). Among the differentially expressed genes in our analysis, all 5 genes that had nuclear/cell division in their functional annotation were downregulated in the asexual species (Sup. Fig. 11). These genes included homologs of MASTL (essential for meiotic division), CDC16 (part of the anaphase promoting complex), two copies of KIF23 (important for the final stage of cell division), and SKA1 (important for chromosome segregation and polar body formation)(36–39). Taken together, these results support the idea that automixis may require subtle but consistent changes in genes involved in cell division.

### The regulation of asexual meiosis in *Artemia*

Two genes found next to each other in the asexuality locus, the calcium channel *ITPR* and the deubiquitinase *USP8*, also showed changes in sequence and/or expression in asexuals, raising the question of whether changes at either are required for asexual reproduction. It is important to note that in the absence of sexual meiosis, some meiotic genes may simply cease to be needed and accumulate disruptive changes neutrally. However, sexual and asexual *Artemia* are only minimally differentiated, and we do not find evidence of meiosis genes located elsewhere in the genome to be systematically disrupted (Sup. Fig. 12). The known functions of these two genes also make them plausible candidates for the switch in reproductive mode. Intracellular calcium, and its regulator *ITPR*, are well established to be key for correct oogenesis progression. In particular, egg activation is triggered by an increase in calcium levels in all species studied so far(40). In many, including mice, this is triggered by fertilization, which activates the inositol triphosphate / *ITPR* pathway and results in the release of the calcium stored in the endoplasmic reticulum(40). *ITPR* is expressed in the *Drosophila* ovary(41), and although it might not be required for the initiation of the calcium wave, it seems to be required for its propagation and release from metaphase I arrest (42). In both Drosophila and several crustaceans, ITPR-mediated activation seems to be independent of fertilization, and rely instead on mechanical or environmental cues (42).

How might modulating ITPR contribute to asexuality under such a scenario? The presence of another functional and expressed copy of ITPR suggests that down-regulation rather than loss of function is at play. One possibility is therefore that reduced expression only allows for a partial resumption of meiosis, and consequent failure to perform a full meiosis I division. To get further insights into this hypothesis, we looked at the top 40 differentially expressed genes in germ cells A for which we could assign a *Drosophila* ortholog, and identified functional clusters among them using StringDB (Sup. Methods, Sup. Fig. 13). Two clusters (with 3 genes each) included genes known to be involved in cytoskeleton organization and cell division, and highlight some downstream processes that may be affected (Figure 5). The up-regulation of two homologs of Tacc, a stabilizer of microtubules (43) and of a homolog of Asp, a gene responsible for spindle pole cohesion (44, 45), combined with the down-regulation of DCX-EMAP, which is associated with the assembly of microtubules (46), suggest that increased stability of microtubules in the meiotic spindle may play a part in disrupting chromosome segregation. The down-regulation of several genes known to be important for cytokinesis (Pav (47), CG33521 (48)) is similarly in line with a reduction of the cytokinesis machinery leading to the skipping of the first meiotic cell division. Consistent with this model, in *Xenopus*, *ITPR* seems to be essential for spindle assembly and emitting the first polar body (49); upon its depletion, polar body emission is reduced, a phenotype reminiscent of skipping the first meiotic division. Interfering with calcium signalling (using strontium chloride for instance) has been used to parthenogenetically activate mouse oocytes, and the blastulation rate increases when cytochalasin B is included in the medium, as it suppresses the polar body extrusion(50, 51). Taken together, these studies suggest that changes at *ITPR* could in principle by themselves facilitate the transition to asexuality, and may have been the first step, consistent with high differentiation of *ITPR2* between sexuals and asexuals.

However, important roles of *USP8* in cell division have also recently been uncovered. Removal of *USP8* in *Drosophila* led to loss of cytoplasmic bridges between cells in ovarian germ cysts, i.e. turned incomplete cell divisions into complete ones (52). *USP8* has also been found to regulate DNA endoreplication in insect salivary gland cells (53). While both of these functions were studied in mitotic cells, *USP8* could play similar roles in regulating DNA replication and cell division in meiosis. In *Drosophila*, *USP8* is also involved in regulating the number of centrosomes (54, 55), the microtubule-organizing centers of cells, something that is crucial for transitions to asexuality (56). While mitotic cells contain one centrosome that gets replicated at each cell division, female meiotic cells degrade theirs, such that embryos rely on centrosomes provided by their fathers at fertilization. Asexuals must therefore either maintain centrosomes through meiosis (57) or recreate them later (56), both of which could be facilitated by changes in *USP8*. While disentangling the potential contributions of *USP8* and *ITPR2* will require genetic manipulation tools that are still being developed in *Artemia*, a likely scenario given their complementary roles in oogenesis and cell division is that downregulation of both genes contributes to asexual meiosis.

### Key pre-adaptations likely facilitated the switch to asexuality

Various studies of parthenogenesis suggest that the modifications that lead to the evolution of asexuality tend to be disruptive changes in meiosis-essential genes or processes, such as the disruptive mutations found at *Rec8* in obligatory parthenogenetic Daphnia. Similarly, hybridization and polyploidy, two scenarios often associated with defects in gametogenesis, are also strongly associated with switches to asexuality(58). In *Artemia*, we observe downregulation of expression and/or disruptive genetic variants at *ITPR* and *USP8*, two genes essential for correct meiotic progression. Some results in mutants of sexual species also support the idea that interfering with different aspects of the meiotic machinery can lead to asexual meiosis. *Drosophila* mutant lines for *yem1*, a gene involved in chromatin remodeling and chromosome behaviour in meiosis I, produce a small number of offspring without paternal genomic contribution(59). Similarly, increasing the copy number of *Polo* (a kinase with meiotic and mitotic functions) and decreasing the expression of *Desat2* (a desaturase) was enough to induce facultative parthenogenesis in *Drosophila melanogaster*(30).

Taken together, these studies show that disturbing different meiotic mechanisms can lead to the emergence of asexual meiosis, making the relative rarity of transitions observed in nature intriguing. One possibility is that new instances of parthenogenesis occur frequently, but are associated with strong deleterious consequences (such as low fertility), and therefore quickly eliminated (60). On the other hand, multiple prerequisite conditions may likely be required before asexuality can arise, limiting the rate at which this occurs (9). In the case of *Artemia*, the acquisition of a new paralog of *ITPR* in Eurasian lineages likely created functional redundancy that facilitated the evolution of asexuality, as *ITPR* is a highly pleiotropic gene. In *Drosophila*, which like other insects only has one copy of *ITPR*, unconditional mutants do not survive past the larval stage(61). A similar scenario may be at play in Daphnia, where multiple paralogs of Rec8 are found. Additionally, it has been suggested that the genetic linkage of different genes regulating the different aspects of meiosis that need to be modified for automixis may be a prerequisite for their coordinated change (9). The birth of *ITPR2* next to *USP8* may have allowed for their coordinated action in asexual meiosis. Finally, low recombination levels in parthenogenetic species with central fusion automixis slows down the loss of heterozygosity that takes place under this form of asexual reproduction, and low recombination levels in the ancestral sexual species may be a prerequisite for the survival of its derived asexual lineages, as has been proposed in *Artemia* (34). The requirement for several pre-adaptations before asexuality can emerge may limit the frequency at which it occurs, which in turn could contribute to the patchy distribution of asexual reproduction throughout the animal tree of life (60). However, the characterizations of minimal steps required for asexual reproduction in other clades is needed to shed light on how general this serendipitous and multi-step acquisition of asexuality is.

## Methods

### DNA extraction and sequencing

For the long-read sequencing, we extracted HMW DNA from a single *Artemia parthenogenetica* (Aibi lake) female and a single *Artemia sinica* female with the Qiagen Genomic-Tip 20/G Kit. The two samples were sequenced on the PromethION nanopore sequencer at the Vienna Biocenter sequencing facility.

For short-read sequencing, we extracted genomic DNA from an individual *Artemia parthenogenetica* female (Atanasovsko) using the Qiagen DNeasy Blood & Tissue kit and sent it to the Vienna BioCenter (VBC) for library-prep and Illumina sequencing.

### Genome Assemblies and Annotation

We downloaded the published male *Artemia sinica* genome assembly (ASM2792156v2, (25)). Since the coding sequence of some genes in this assembly contained premature stop codons or frameshifts, we polished it using Hapo-G(62) with short genomic read data from a published *A. sinica* male individual (SRR19741769). We ran Merqury(63) to verify the improvement in the assembly quality (Sup. Table 4). For the annotation, we used HISAT2(64) to align the available *Artemia sinica* female gonad samples (SRR15446679, SRR15446678, SRR15446677, SRR15446676) to the genome and used StringTie2(65) to generate a GTF with the gene models. For the *Artemia parthenogenetica* genome we assembled the ONT long read (see section DNA extraction and sequencing) using Flye(66) with the following options: “--nano-hq --genome-size 1g --scaffold”. We used the same long reads to purge the duplicated sequences with minimum alignment score of 90 in the assembly using purge_dups(67). We scaffolded the assembly using RagTag(68) with the 21 chromosomes from the polished *Artemia sinica* genome as the reference. We re-assembled the genome of *Artemia sp. Kazakhstan* from published male short read genomic data (SRR19741746, SRR19741779)(16, 25, 69) using MEGAHIT(70) and scaffolded the genomes with the same reads using SOAPdenovo-fusion(68). The generated scaffolds were anchored with RagTag(68) with the *Artemia sinica* chromosome-level genome assembly as the reference (ASM2792156v1).

### Nuclei isolation, single-nucleus RNA sequencing and analysis

Adult *A. parthenogenetica* (from a stock originally from Aibi Lake in China) and *A. sp. Kazakhstan* were collected from colonies maintained in the lab. We washed adult *Artemia* females in MilliQ water for 15 minutes and then dissected the female reproductive organs under a stereomicroscope, collected them in PBS, and placed them on ice. After dissecting, we followed the same protocol described in(19, 71) to isolate nuclei. The samples were sent to the Vienna BioCenter sequencing facility for sorting, library preparation and 10x 3’ GEX single-cell sequencing. We generated four samples with 25 females each: two from *A. parthenogenetica* and two from *A. sp. Kazakhstan*.

The single-nucleus RNAseq reads were aligned to the polished *A. sinica* genome using CellRanger(72) with the newly generated GTF (see section Genome Assemblies and Annotation). The resulting count matrices were corrected using CellBender (73) to eliminate technical artifacts. The corrected matrices were then loaded into Seurat (22) and filtered for nCount_RNA >250. The detection of the highly variable genes was performed using DUBStepR(74), and the integration of the different replicates was performed using Harmony(75). For the cell type annotation, we used SAMap (20) to map the single nucleus atlas generated in this study to the *Artemia franciscana* snRNAseq atlas (19) and transferred the labels based on the alignment scores. The number of nuclei per cell type recovered for each replicate is provided in Sup. table 5).

For the differential expression analysis between the sexual and asexual species, we used two complementary approaches to minimize the number of false positives. The first was to aggregate the expression counts for each gene within clusters, which resulted in two replicates per species (pseudobulks), and then use DESeq2 (21) to test for differential expression. The second approach used the Seurat FindMarkers function to find the differentially expressed genes between all the cells of one cluster in one species against the other. In this approach, MAST(23) was used for differential testing and the replicates were used as a latent variable.

### Backcrossing experiment, F_ST_ analysis and chromosome painting

We mated three *A. parthenogenetica* rare males (from a colony originated from a stock from Lake Atanasovsko, Bulgaria) with unmated females from *Artemia sp. Kazakhstan*. The resulting F1 offspring were split into vials before the juvenile stage, and 8 F1 males were collected to backcross to *A. sp. Kazakh* unmated females. The F2 offspring were isolated into individual vials, and 9 females which reproduced asexually were frozen instantly upon the appearance of nauplii. After three months, 9 of the remaining females, which did not produce any offspring, were frozen as control females. Genomic DNA was extracted from the 18 samples using the Qiagen DNeasy Blood & Tissue kit and sent to the Vienna BioCenter (VBC) for Illumina short read sequencing. One of the control samples did not have enough material for sequencing; therefore, only 17 individuals were included in the analysis.

We aligned the genomic reads of the 17 individuals from the newly generated backcrossing experiment and the previously published data from the 15 F2 female individuals from a *A. parthenogenetica* rare male x *A. sp. Kazakhstan* male backcross (SRR19741752 to SRR19741757 and SRR19741759 to SRR19741767) (25) to the *Artemia sinica* genome using segemehl(76). We called SNPs using BCFtools(77) for all the samples along with genomic samples for two females of *A. sp. Kazakhstan* (SRR19741778 and SRR19741777), one female *A. parthenogenetica* (Aibi lake, SRR19741775), and one newly generated female sample from *Artemia parthenogenetica* (Atanasovsko, see section DNA extraction and sequencing). After the SNP calling, F_ST_ between the asexual and control female populations (*A. sp. Kaz* and *A. par* samples were removed from the vcf before estimating F_ST_) was calculated using vcftools (--weir-fst-pop)(78). The negative F_ST_ values were clipped to 0 and the rolling median of 100 consecutive positions was used to generate the F_ST_ plots (Figure 2 A and B).

The SNPs from the 2 *A. sp Kazakhstan* females and 2 asexual females were used to identify variants fixed in the parental species, in order to use them for identifying the patterns of inheritance of genomic regions. We filtered for SNPs that had the following genotypes: 0/0 in asexuals and 1/1 in *Artemia sp. Kazakhstan* or 1/1 in asexuals and 0/0 *Artemia sp. Kazakhstan*). Genomic windows of 2MB in each individual were assigned *A. parthenogenetica* ancestry if 60% or more of the SNPs were either homozygous or heterozygous for the *A. par* allele, and they were assigned *A. sp. Kaz* ancestry if 60% of the SNPs or more were homozygous for the *A. sp. Kaz* allele. Windows that did not fit within those categories were left unassigned.

### SnpEff analysis

We filtered the SNP calls from the 2 *Artemia sp. Kazakhstan* and 2 *A. parthenogenetica* individuals (described in the previous section) for SNPs that are homozygous in *A. sp. Kazakhstan* for the reference allele (0/0) and homozygous or heterozygous for the alternative allele (0/1 or 1/1) in *Artemia parthenogenetica*. We then used SnpEff (26) to predict the variant effects. The coding sequences and protein predictions for building the database were generated using TransDecoder (https://github.com/TransDecoder/TransDecoder/wiki). Only one isoform per gene was included, and for transcripts that had multiple peptide predictions in the same transcript, the sum of the variants is shown in Figure 3.

### Structural variant calling and variant effect prediction

We mapped the ONT long reads from *Artemia parthenogenetica* and *Artemia sinica* (see section DNA extraction and sequencing) to the polished *Artemia sinica* genome using minimap2(79) with the following options: “-a -m 5 -s 5 -n 1 -B2 -O2,24 -E2,1”. We removed the secondary alignments using (samtools view -F0×100). We used cuteSV to call structural variants in the *A. parthenogenetica* sample, filtered for variants with more than two supporting reads, and then genotyped the two samples for the identified SVs. We filtered for variants with 0/0 in *A. sinica* and either 0/1 or 1/1 in *A. parthenogenetica*. We used VEP (28) to predict the variant effects.

### Identification of protein coding gene in the putative asexuality locus

The protein coding genes were identified by running LncDC(80) to assign a biotype (either mrna or lncRNA) for all transcripts. The transcripts in the putative asexuality region (Figure 3 D, Sup. Table 2) were annotated by blasting the protein coding genes to the annotated *A. franciscana* proteome on NCBI(81).

### *ITPR* phylogenetic analysis

To explore the phylogenetic relationship between *A. franciscana*’s two copies of ITPR (annotated as ITPR and ITPR-like) and the *A. sinica* ones, we mapped the protein sequence of the *A. fanciscana* ITPR using miniprot(82) to the genomes of the fairy shrimp *Branchinecta sandiegonensis* (NCBI accession GCA_030848855.1), *Artemia sinica*(25), *Artemia parthenogenetica* (produced in this study from long reads, see section Genome Assemblies and Annotation), and *Artemia sp. Kazakhstan* (produced in this study using short reads, see Genome Assemblies and Annotation). To produce the phylogenetic trees, we extracted the miniport annotated nucleotide sequences from all the genomes, aligned them, and generated the trees using maximum likelihood (PhyML) on NGphylogeny.fr(83).

### Bulk RNAseq analysis

We mapped the female gonad RNAseq samples (Sup. table 6) generated in (16) to the polished *Artemia sinica genome* using HISAT2(64). We used StringTie2(65) to generate the counts per gene for each sample, and DEseq2 (21) to perform differential expression analysis between the sexual and asexual individuals of the 6 species.

## Data availability statement

The scripts used in the analysis are available on the GitHub page: https://github.com/Melkrewi/Artemia_parthenogenesis. The raw data is available on the NCBI short read archive (BioProject number XXXXXX). The Seurat objects, differential expression analysis results, and VCF files are available for download on ISTA Research Explorer (ISTA REx) using the following link: XXXXX. The single nucleus atlas can be viewed with this link on the UCSC Cell Browser: XXXXX. [All “XXXXXX”s will be replaced with URLs/accession numbers before publication]. The *Artemia sinica* proteome annotated using StringDB (24) is accessible using the following link: https://version-12-0.string-db.org/organism/STRG0A32TIE.

## References

1. S. P. Otto, The Evolutionary Enigma of Sex. Am. Nat. 174, S1–S14 (2009).

2. A. Brandt, et al., Haplotype divergence supports long-term asexuality in the oribatid mite Oppiella nova. Proc. Natl. Acad. Sci. U. S. A. 118, e2101485118 (2021).

3. C. J. van der Kooi, C. Matthey-Doret, T. Schwander, Evolution and comparative ecology of parthenogenesis in haplodiploid arthropods. Evol. Lett. 1, 304–316 (2017).

4. G. Esposito, et al., First report of recurrent parthenogenesis as an adaptive reproductive strategy in the endangered common smooth-hound shark Mustelus mustelus. Sci. Rep. 14, 17171 (2024).

5. J. Engelstädter, Asexual but Not Clonal: Evolutionary Processes in Automictic Populations. Genetics 206, 993–1009 (2017).

6. B. D. Eads, D. Tsuchiya, J. Andrews, M. Lynch, M. E. Zolan, The spread of a transposon insertion in Rec8 is associated with obligate asexuality in Daphnia. Proc. Natl. Acad. Sci. 109, 858–863 (2012).

7. J. Jaquiéry, et al., Genetic Control of Contagious Asexuality in the Pea Aphid. PLOS Genet. 10, e1004838 (2014).

8. B. Yagound, et al., A Single Gene Causes Thelytokous Parthenogenesis, the Defining Feature of the Cape Honeybee *Apis mellifera capensis*. Curr. Biol. 30, 2248–2259.e6 (2020).

9. M. Neiman, T. F. Sharbel, T. Schwander, Genetic causes of transitions from sexual reproduction to asexuality in plants and animals. J. Evol. Biol. 27, 1346–1359 (2014).

10. B. B. Normark, L. R. Kirkendall, “Chapter 192 - Parthenogenesis in Insects and Mites” in Encyclopedia of Insects (Second *Edition)*, V. H. Resh, R. T. Cardé, Eds. (Academic Press, 2009), pp. 753–757.

11. O. Nougué, et al., Automixis in Artemia: solving a century-old controversy. J. Evol. Biol. 28, 2337–2348 (2015).

12. L.-Y. Xu, et al., A cytological revisit on parthenogenetic Artemia and the deficiency of a meiosis-specific recombinase DMC1 in the possible transition from bisexuality to parthenogenesis. Chromosoma 132, 89–103 (2023).

13. S. Kumar, G. Stecher, M. Suleski, S. B. Hedges, TimeTree: A Resource for Timelines, Timetrees, and Divergence Times. Mol. Biol. Evol. 34, 1812–1819 (2017).

14. A. Eimanifar, G. Van Stappen, M. Wink, Geographical distribution and evolutionary divergence times of Asian populations of the brine shrimp Artemia (Crustacea, Anostraca). Zool. J. Linn. Soc. 174, 447–458 (2015).

15. T. H. Oakley, J. M. Wolfe, A. R. Lindgren, A. K. Zaharoff, Phylotranscriptomics to Bring the Understudied into the Fold: Monophyletic Ostracoda, Fossil Placement, and Pancrustacean Phylogeny. Mol. Biol. Evol. 30, 215–233 (2013).

16. A. K. Huylmans, A. Macon, F. Hontoria, B. Vicoso, Transitions to asexuality and evolution of gene expression in Artemia brine shrimp. Proc. Biol. Sci. 288, 20211720 (2021).

17. M. Maccari, A. Gómez, F. Hontoria, F. Amat, Functional rare males in diploid parthenogenetic Artemia. J. Evol. Biol. 26, 1934–1948 (2013).

18. M. Maccari, F. Amat, F. Hontoria, A. Gómez, Laboratory generation of new parthenogenetic lineages supports contagious parthenogenesis in Artemia. PeerJ 2, e439 (2014).

19. M. Elkrewi, B. Vicoso, Single-nucleus atlas of the Artemia female reproductive system suggests germline repression of the Z chromosome. PLOS Genet. 20, e1011376 (2024).

20. A. J. Tarashansky, et al., Mapping single-cell atlases throughout Metazoa unravels cell type evolution. eLife (2021). Available at: https://elifesciences.org/articles/66747 [Accessed 5 February 2024].

21. M. I. Love, W. Huber, S. Anders, Moderated estimation of fold change and dispersion for RNA-seq data with DESeq2. Genome Biol. 15, 550 (2014).

22. Y. Hao, et al., Dictionary learning for integrative, multimodal and scalable single-cell analysis. Nat. Biotechnol. 1–12 (2023). 10.1038/s41587-023-01767-y.

23. G. Finak, et al., MAST: a flexible statistical framework for assessing transcriptional changes and characterizing heterogeneity in single-cell RNA sequencing data. Genome Biol. 16, 278 (2015).

24. S. D, et al., STRING v10: protein-protein interaction networks, integrated over the tree of life. Nucleic Acids Res. 43 (2015).

25. M. Elkrewi, et al., ZW sex-chromosome evolution and contagious parthenogenesis in Artemia brine shrimp. Genetics 222, iyac123 (2022).

26. P. Cingolani, et al., A program for annotating and predicting the effects of single nucleotide polymorphisms, SnpEff. Fly (Austin) 6, 80–92 (2012).

27. T. Jiang, et al., Long-read-based human genomic structural variation detection with cuteSV. Genome Biol. 21, 189 (2020).

28. W. McLaren, et al., The Ensembl Variant Effect Predictor. Genome Biol. 17, 122 (2016).

29. T. Rausch, et al., DELLY: structural variant discovery by integrated paired-end and split-read analysis. Bioinformatics 28, i333–i339 (2012).

30. A. L. Sperling, D. K. Fabian, E. Garrison, D. M. Glover, A genetic basis for facultative parthenogenesis in Drosophila. Curr. Biol. CB 33, 3545–3560.e13 (2023).

31. A. N. Beachum, et al., Orphan nuclear receptor ftz-f1 (NR5A3) promotes egg chamber survival in the Drosophila ovary. G3 GenesGenomesGenetics 11, jkab003 (2021).

32. E. Seronick, et al., CRISPR/Cas9 genome editing system confirms centriolin’s role in cytokinesis. BMC Res. Notes 15, 8 (2022).

33. A. Grallert, et al., A PP1/PP2A phosphatase relay controls mitotic progression. Nature 517, 94–98 (2015).

34. N. O. Rode, et al., The Origin of Asexual Brine Shrimps. Am. Nat. 200, E52–E76 (2022).

35. C. R. Haag, L. Theodosiou, R. Zahab, T. Lenormand, Low recombination rates in sexual species and sex–asex transitions. Philos. Trans. R. Soc. B Biol. Sci. 372, 20160461 (2017).

36. J. A. Pesin, T. L. Orr-Weaver, Developmental Role and Regulation of cortex, a Meiosis-Specific Anaphase-Promoting Complex/Cyclosome Activator. PLOS Genet. 3, e202 (2007).

37. N. J. Camlin, E. A. McLaughlin, J. E. Holt, Motoring through: the role of kinesin superfamily proteins in female meiosis. Hum. Reprod. Update 23, 409–420 (2017).

38. Q.-H. Zhang, et al., Localization and function of the Ska complex during mouse oocyte meiotic maturation. Cell Cycle Georget. Tex 11, 909–916 (2012).

39. M.-Y. Kim, et al., Bypassing the Greatwall–Endosulfine Pathway: Plasticity of a Pivotal Cell-Cycle Regulatory Module in Drosophila melanogaster and Caenorhabditis elegans. Genetics 191, 1181–1197 (2012).

40. B. Lee, S.-Y. Yoon, C. Malcuit, J. B. Parys, R. A. Fissore, Inositol 1,4,5-Trisphosphate Receptor 1 Degradation in Mouse Eggs and Impact on [Ca2+]i Oscillations. J. Cell. Physiol. 222, 238–247 (2010).

41. C. V. Sartain, M. F. Wolfner, Calcium and egg activation in Drosophila. Cell Calcium 53, 10–15 (2013).

42. T. Kaneuchi, et al., Calcium waves occur as Drosophila oocytes activate. Proc. Natl. Acad. Sci. 112, 791–796 (2015).

43. F. Gergely, D. Kidd, K. Jeffers, J. G. Wakefield, J. W. Raff, D–TACC: a novel centrosomal protein required for normal spindle function in the early Drosophila embryo. EMBO J. 19, 241–252 (2000).

44. A. Ito, G. Goshima, Microcephaly protein Asp focuses the minus ends of spindle microtubules at the pole and within the spindle. J. Cell Biol. 211, 999–1009 (2015).

45. M. G. Riparbelli, M. Migliorini, G. Callaini, Astral Microtubules Are Dispensable for Pavarotti Localization During Drosophila Spermatogonial Mitoses. Cytoskeleton 82, 516–528 (2025).

46. X. Song, et al., DCX-EMAP is a core organizer for the ultrastructure of Drosophila mechanosensory organelles. J. Cell Biol. 222, e202209116 (2023).

47. C. Cabernard, Cytokinesis in Drosophila melanogaster. Cytoskeleton 69, 791–809 (2012).

48. D. Fagegaltier, et al., Oncogenic transformation of Drosophila somatic cells induces a functional piRNA pathway. Genes Dev. 30, 1623–1635 (2016).

49. R. Li, et al., Inositol 1, 4, 5-trisphosphate receptor is required for spindle assembly in Xenopus oocytes. Mol. Biol. Cell 33, br27 (2022).

50. K. Versieren, B. Heindryckx, S. Lierman, J. Gerris, P. D. Sutter, Developmental competence of parthenogenetic mouse and human embryos after chemical or electrical activation. Reprod. Biomed. Online 21, 769–775 (2010).

51. S.-F. Ma, et al., Parthenogenetic activation of mouse oocytes by strontium chloride: A search for the best conditions. Theriogenology 64, 1142–1157 (2005).

52. J. Mathieu, P. Michel-Hissier, V. Boucherit, J.-R. Huynh, The deubiquitinase USP8 targets ESCRT-III to promote incomplete cell division. Science 376, 818–823 (2022).

53. W. Qian, et al., USP8 and Hsp70 regulate endoreplication by synergistically promoting Fzr deubiquitination and stabilization. Sci. Adv. 11, eadq9111 (2025).

54. M. Kwon, et al., Mechanisms to suppress multipolar divisions in cancer cells with extra centrosomes. Genes Dev. 22, 2189–2203 (2008).

55. S. Darling, A. B. Fielding, D. Sabat-Pośpiech, I. A. Prior, J. M. Coulson, Regulation of the cell cycle and centrosome biology by deubiquitylases. Biochem. Soc. Trans. 45, 1125–1136 (2017).

56. M. G. Riparbelli, M. Gottardo, G. Callaini, Parthenogenesis in Insects: The Centriole Renaissance. Results Probl. Cell Differ. 63, 435–479 (2017).

57. A. Perrier, et al., Maternal inheritance of functional centrioles in two parthenogenetic nematodes. Nat. Commun. 15, 6042 (2024).

58. S. Freitas, et al., Parthenogenesis in Darevskia lizards: A rare outcome of common hybridization, not a common outcome of rare hybridization. Evolution 76, 899–914 (2022).

59. R. E. Meyer, M. Delaage, R. Rosset, M. Capri, O. Aït-Ahmed, A single mutation results in diploid gamete formation and parthenogenesis in a Drosophila yemanuclein-alpha meiosis I defective mutant. BMC Genet. 11, 104 (2010).

60. T. Schwander, B. J. Crespi, Twigs on the tree of life? Neutral and selective models for integrating macroevolutionary patterns with microevolutionary processes in the analysis of asexuality. Mol. Ecol. 18, 28–42 (2009).

61. K. Venkatesh, G. Siddhartha, R. Joshi, S. Patel, G. Hasan, Interactions between the inositol 1,4,5-trisphosphate and cyclic AMP signaling pathways regulate larval molting in Drosophila. Genetics 158, 309–318 (2001).

62. J.-M. Aury, B. Istace, Hapo-G, haplotype-aware polishing of genome assemblies with accurate reads. NAR Genomics Bioinforma. 3, lqab034 (2021).

63. A. Rhie, B. P. Walenz, S. Koren, A. M. Phillippy, Merqury: reference-free quality, completeness, and phasing assessment for genome assemblies. Genome Biol. 21, 245 (2020).

64. D. Kim, J. M. Paggi, C. Park, C. Bennett, S. L. Salzberg, Graph-based genome alignment and genotyping with HISAT2 and HISAT-genotype. Nat. Biotechnol. 37, 907–915 (2019).

65. S. Kovaka, et al., Transcriptome assembly from long-read RNA-seq alignments with StringTie2. Genome Biol. 20, 278 (2019).

66. M. Kolmogorov, J. Yuan, Y. Lin, P. A. Pevzner, Assembly of long, error-prone reads using repeat graphs. Nat. Biotechnol. 37, 540–546 (2019).

67. D. Guan, et al., Identifying and removing haplotypic duplication in primary genome assemblies. Bioinformatics 36, 2896–2898 (2020).

68. M. Alonge, et al., Automated assembly scaffolding using RagTag elevates a new tomato system for high-throughput genome editing. Genome Biol. 23, 258 (2022).

69. A. K. Huylmans, M. A. Toups, A. Macon, W. J. Gammerdinger, B. Vicoso, Sex-Biased Gene Expression and Dosage Compensation on the *Artemia franciscana* Z-Chromosome. Genome Biol. Evol. 11, 1033–1044 (2019).

70. D. Li, C.-M. Liu, R. Luo, K. Sadakane, T.-W. Lam, MEGAHIT: an ultra-fast single-node solution for large and complex metagenomics assembly via succinct de Bruijn graph. Bioinformatics 31, 1674–1676 (2015).

71. C. N. McLaughlin, Y. Qi, S. R. Quake, L. Luo, H. Li, Isolation and RNA sequencing of single nuclei from Drosophila tissues. STAR Protoc. 3, 101417 (2022).

72. G. X. Y. Zheng, et al., Massively parallel digital transcriptional profiling of single cells. Nat. Commun. 8, 14049 (2017).

73. S. J. Fleming, et al., Unsupervised removal of systematic background noise from droplet-based single-cell experiments using CellBender. Nat. Methods 20, 1323–1335 (2023).

74. B. Ranjan, et al., DUBStepR is a scalable correlation-based feature selection method for accurately clustering single-cell data. Nat. Commun. 12, 1–12 (2021).

75. I. Korsunsky, et al., Fast, sensitive and accurate integration of single-cell data with Harmony. Nat. Methods 16, 1289–1296 (2019).

76. C. Otto, P. F. Stadler, S. Hoffmann, Lacking alignments? The next-generation sequencing mapper segemehl revisited. Bioinformatics 30, 1837–1843 (2014).

77. P. Danecek, et al., Twelve years of SAMtools and BCFtools. GigaScience 10, giab008 (2021).

78. P. Danecek, et al., The variant call format and VCFtools. Bioinformatics 27, 2156–2158 (2011).

79. H. Li, Minimap2: pairwise alignment for nucleotide sequences. Bioinformatics 34, 3094–3100 (2018).

80. M. Li, C. Liang, LncDC: a machine learning-based tool for long non-coding RNA detection from RNA-Seq data. Sci. Rep. 12, 19083 (2022).

81. E. Jo, et al., High-quality chromosome-level genome assembly of female *Artemia franciscana* reveals sex chromosome and *Hox* gene organization. Heliyon 10, e38687 (2024).

82. H. Li, Protein-to-genome alignment with miniprot. Bioinformatics 39, btad014 (2023).

83. F. Lemoine, et al., NGPhylogeny.fr: new generation phylogenetic services for non-specialists. Nucleic Acids Res. 47, W260–W265 (2019).

